# Environmental and Socioeconomic Factors Associated with West Nile Virus

**DOI:** 10.1101/349720

**Authors:** E Hernandez, AL Joyce, R Torres

## Abstract

Environmental and socioeconomic risk factors associated with West Nile Virus cases were investigated in the Northern San Joaquin Valley region of California, a largely rural area. The study included human West Nile Virus (WNV) cases from the years 2011-2015 in the three county area of San Joaquin, Stanislaus and Merced Counties, and examined whether factors were associated with WNV using census tracts as the unit of analysis. Environmental factors included temperature, precipitation, mosquitoes positive for WNV, and habitat. Socioeconomic variables included age, education, housing age, home vacancies, median income, population density, ethnicity, and language spoken. Chi-squared independence tests were used to examine whether each variable was associated with WNV in each county, and then also used for the three counties combined. Logistic regression was used for a three-county combined analysis, to examine which environmental and socioeconomic variables were most likely associated with WNV cases. The chi-squared tests found that the variables associated with WNV varied in each of the three counties. The chi-squared tests for data combined from the three counties found that WNV cases were significantly associated with mosquitoes positive for WNV, urban habitat, higher home vacancies, higher population density, higher education, and ethnicity. Logistic regression analysis revealed that overall, the environmental factors precipitation, mean temperature, and WNV positive mosquitoes were the strongest predictors of WNV cases. Results support efforts of mosquito control districts, which aim for source reduction of mosquito breeding sites. In addition, findings suggest that residents with higher income and education may be more aware of WNV and its symptoms, and more likely to request testing from physicians. Lower income and education residents may not be aware of WNV. Public health education might increase its prevention messages about vector-borne disease in the various languages of the region, which would contribute overall to public health in the region.

## Introduction

Mosquito-borne illnesses are a threat to global health and are known to affect nearly 700 million people annually (1). One of these diseases is West Nile Virus (WNV), an arthropod-borne virus of the family *Flaviviridae* that is commonly spread by mosquitoes (2). This virus emerged from Uganda in 1937 and since this initial discovery, has spread geographically to regions including Europe, Australia, Asia, the Caribbean, and South America (3). In 1999, WNV reached the United States when its first reported case emerged in New York City with a group of patients diagnosed with encephalitis (4). By July 2003, WNV arrived in California and shortly afterwards its activity was detected across all 58 counties (5).

WNV is maintained through a cycle between birds and mosquitoes. It is transmitted primarily by mosquitoes that belong to the genus *Culex: Culex pipiens, Culex quinquefasciatus,* and *Culex tarsalis* are the main carriers (6). A small proportion of human infections can develop from blood transfusions, organ transplants, and transmittance from mother to child during pregnancy, delivery, or through breastfeeding (7). WNV is considered the arthropod-borne disease responsible for the greatest number of neuroinvasive disease outbreaks that have ever been reported (8). However, many humans who are infected are asymptomatic or experience minor symptoms (9). People who are 50 years of age or older are at the greatest risk of developing severe illnesses (10). Currently, no vaccine exists for humans; treatment for mild cases such as over-the-counter pain relievers to reduce fevers or joint pains are available.

The importance of environmental factors and their influence of where human infections occur have been investigated since WNV arrived to the United States (11). Several variables that previous studies found associated with WNV include temperature, rainfall, and habitat. In southern California, summer mean temperature, land surface temperature, elevation, landscape diversity, and vegetation water content were principle environmental factors that contributed to WNV propagation (12). High temperature has been consistently associated with outbreaks of WNV (13). Above-average summer temperatures were closely linked to hot spots of WNV activity in the United States from 2002-2004 (14). Specific habitats also permit species of mosquitoes to thrive. In Shelby County, Tennessee, *Cx. pipiens* and *Cx. quinquefasciatus* were positively associated with urban habitats (15). The abundance of *Cx. pipiens* complex mosquitoes however, was lower in rural sites. Above-average precipitation may also lead to greater mosquito abundance and an increase for WNV outbreaks in humans (16,17), but in urban areas, a prolonged time of drought can result in increased abundance of mosquitoes that can intensify transmission of WNV (13).

Socioeconomic variables and anthropogenic characteristics of the environment also contribute to predicting WNV prevalence in several studies. Areas with lower per capita income in Orange County had higher prevalence levels of WNV in vectors (18). The density of neglected swimming pools associated with home foreclosures provided an explanation for years of high WNV prevalence in this area (19). Housing unit density, neglected swimming pools, mean per capita income, increased mosquito breeding sites and ditches, and older housing average age were additional risk factors for Orange County, California (20). In Suffolk County, New York urbanization and increased WNV activity were associated; fragmented natural areas, increased road density, and urban areas where there were high numbers of people with a college education had higher levels of WNV (21).

This study investigated environmental and socioeconomic factors associated with human WNV cases in the Northern San Joaquin Valley region of the Central Valley of California. The region is largely rural and comprised of three counties, each with several moderate-sized cities. Variables previously investigated in other studies were included, as were several new variables. Environmental variables herein consisted of precipitation, temperature, mosquitoes which tested positive for WNV, and habitat. Socioeconomic variables included age, education, housing age, home vacancies, income, and population density. Two new variables were considered which were not examined in previous studies; ethnicity and language spoken at home. These two variables were also considered to investigate whether non-white ethnic groups or non-English speakers may be more likely to contract WNV due to language barriers or difficulty in accessing media and educational materials about WNV prevention. The objective was to determine whether the presence of human WNV cases in each census tract was associated with these environmental or socioeconomic variables, in order to examine where most human WNV cases occurred in the region.

## Methods

### Study area

The areas of study consisted of three counties in the Northern San Joaquin Valley of California. From north to south, the three counties were San Joaquin, Stanislaus, and Merced County (Fig 1). In this region, there are two mosquito species, *Cx. pipiens* and *Cx. tarsalis*, which are the primary vectors of WNV (22).

**Fig 1.**
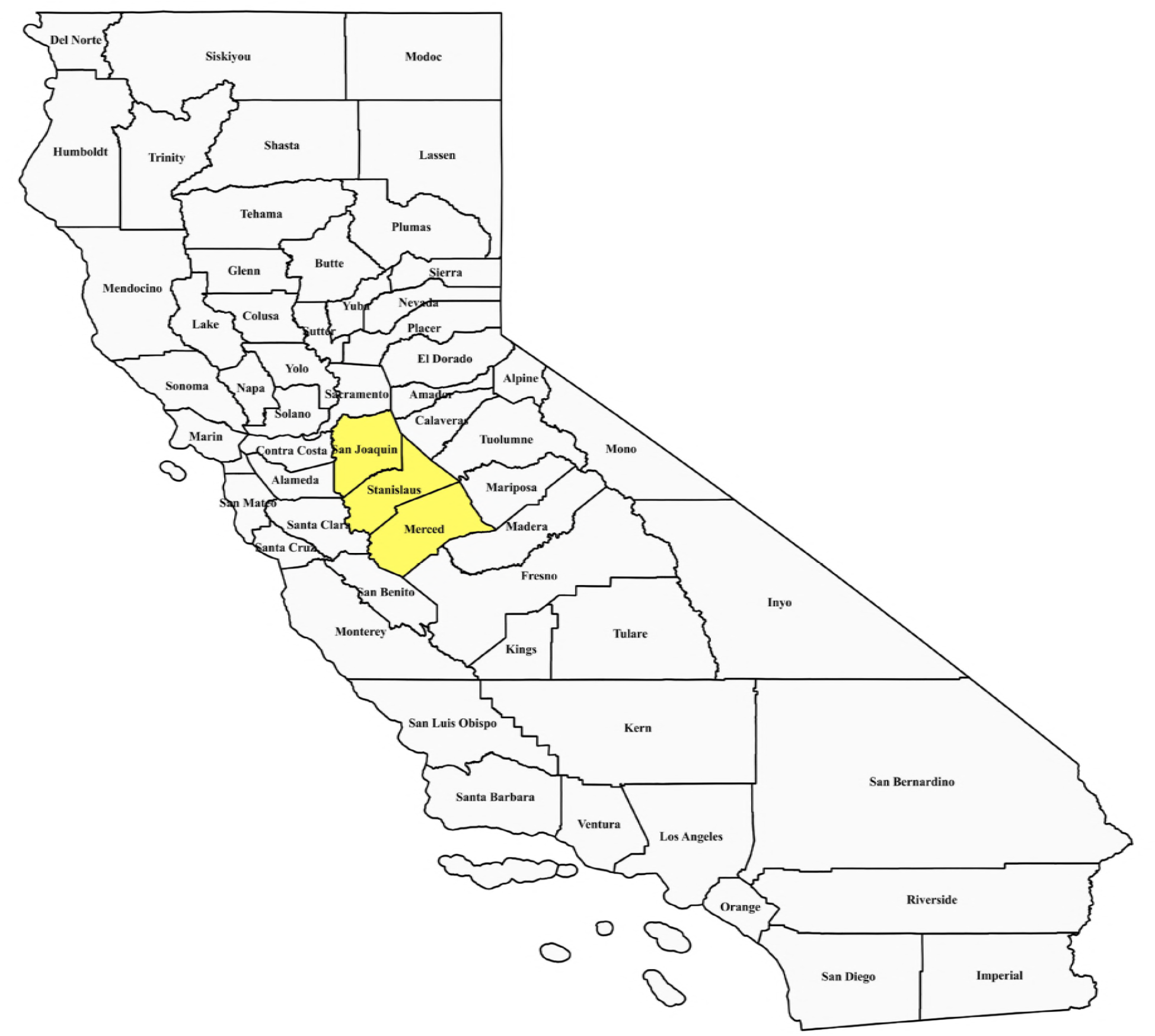
Map of California counties. Three counties in the Northern San Joaquin valley were included in the study. From the north to south, the three counties are San Joaquin, Stanislaus, and Merced (all shown in yellow). The map was created from census tract layers retrieved from the 2010 Census TIGER/Line files using software QGIS 2.18.9. Census tract layers and software can be accessed at https://www.census.gov/cgi-bin/geo/shapefiles2010/main and https://qgis.org/en/site/.

San Joaquin County is located approximately 91 miles east of San Francisco and just south of the Sacramento area. According to the 2010 United States Census, the population size of San Joaquin County was 685,306 and had a total area of 1,426 square miles (23). Large cities within this county with over 50,000 people are Manteca, Stockton, Lodi, and Tracy. In San Joaquin County, 76.7% of people had a high school diploma or higher while 17.5% had a bachelor’s degree or higher. The median household income from the 2010 U.S. Census was $54,341. The county is a rural area and its ethnic composition in 2010 was 51.0% White, 25.9% Hispanic, 14.4% Asian, 7.6% African American, and 1.1% American Indian.

Stanislaus County is located south of San Joaquin County and north of Merced County (Fig 1). The population size in the county in 2010 was 514,453, and the county has a total area of 1,514 square miles. Stanislaus County contains three or four cities with over 50,000 people each, which include Modesto and Turlock. In this county, 75.3% of the population had a high school diploma or higher while 16.3% had a bachelor’s degree or higher. The county’s median household income from the 2010 Census was $51,094. The ethnic makeup of this county consisted of 65.6% White, 25.3% Hispanic, 5.1% Asian, 2.9% African American, and 1.1% American Indian.

Merced County is at the southern end of the three counties. As of the 2010 United States Census, the population size was 255,793. The total area of the county is approximately 1,978 square miles. Merced is the only city with over 50,000 people; the next two largest cities are Los Banos (35,972) and Atwater (28,168) (23). Approximately 67.0% of people living in Merced County had a high school diploma or higher while 12.5% had a bachelor’s degree or higher. The median household income from the 2010 Census was $43,844. The ethnic composition of Merced County was 58.0% White, 29.3% Hispanic, 7.4% Asian, 3.9% African American, and 1. 4% American Indian.

### Data collection

Data used in this study include environmental and socioeconomic variables, and the number of human West Nile Virus (WNV) cases from 2011-2015 in each census tract of San Joaquin, Stanislaus, and Merced Counties. The environmental variables include precipitation, temperature, the number of mosquitoes positive for WNV, and the habitat for each census tract. Annual precipitation and temperature data for the year 2011 (minimum, mean, and maximum temperature) were acquired from the Oregon State University PRISM Climate Group (24). Using the PRISM data, the annual mean precipitation, and mean high and mean low temperatures were determined for a central location in each census tract of the study. The latitude and longitude from each collection site for each WNV positive mosquito was used to determine the census tract where the mosquito was collected. The location of each WNV positive mosquito in the three counties for years 2011-2015 was obtained from the California Vectorborne Disease Surveillance System (CALSURV), a database for mosquito management in California (25). Only *Cx. pipiens* and *Cx. tarsalis* mosquitoes which tested positive for WNV and which were included in CALSURV were used for analyses. To categorize census tracts by habitat, Google Earth was used to view each tract and classify it as rural or non-rural. The definition of rural was used from the US Census Bureau, which was ‘census tracts with less than 2500 people per square mile’ (26).

Socioeconomic variables were obtained from the 2010 United States Census Bureau Census. Socioeconomic variables were determined for each of the 281 census tracts used from the three counties, and include the following: age, education, housing age, home vacancies, median income, population density, ethnicity, and language spoken (S1 Table). For the variable age, the percent of the population over age 65 was used for each census tract. For education, the percent of the population with a high school education or higher was used. The median year homes were constructed in each census tract was used for housing age. The number of housing vacancies for each tract was determined, using the US Census ‘housing vacancies-other’ category. For ethnicity, the percent white was included. The variable language spoken was defined as the percent of households in a census tract speaking English. The number of human WNV cases in each census tract in San Joaquin, Stanislaus, and Merced Counties during 2011-2015 was obtained from the California Department of Public Health Infectious Diseases Branch, Surveillance and Statistics Section. All WNV cases were completely anonymous, and only identified as a case at the census track level. Human subjects approval was obtained from Institutional Review Board (IRB) at University of California Merced and from the California Department of Public Health.

## Statistical analysis

### Descriptive overview of human West Nile cases in census tracts

The number of human WNV cases which occurred from 2011-2015 were determined for each of the three counties. Subsequently, the number of census tracts with and without human WNV cases were determined for each county. One census tract in San Joaquin County was dropped from analyses due to incomplete data. For each year from 2011-2015, the frequency of human WNV cases in each county were plotted for comparison. For each county, a map was made using the software QGIS 2.18.9 (27). Census tract layers for each county were downloaded from the 2010 Census TIGER/Line files (28). This information was used to illustrate census tracts with and without human WNV cases. In addition, a map was created that contains all positive human WNV cases and WNV positive mosquito cases plotted in each census tract for each county. All data were summarized and analyzed at the census tract level.

### Association tests for Factors and Human WNV cases

Chi-square independence tests were run to determine whether a significant association existed between each factor and human WNV cases. Tests were first run for each county, and then for the three counties combined. Variables tested were West Nile positive mosquitoes, habitat, age, education, income, housing age, housing vacancies, population density, ethnicity, and language. For each variable (for example age), we used the 2010 Census to first determine the mean for the variable for each county. For example, for age, the mean percent of the population over age 65 was used to then categorize the census tracts in the county into those with a ‘lower’ or ‘higher’ than the average percent of the population over age 65. Subsequently, a chi-square independence test was used to determine if WNV human cases were associated with age. A similar analysis was run for the other ten variables, by first finding the mean for the variable for the county, and then finding census tracts for each variable which were ‘lower’ and ‘higher’ than average to run the chi-square test. For two variables, income and housing year, the US Census provided median values, and the analyses used the median values rather than mean. STATA 14.2 was used to run these analyses (29).

A chi-square independence test was also run for each variable with the combined data from the three counties. For most of the ten variables, the mean for the category was found by averaging the mean for that variable from the three counties; census tracts were then again classified into ‘low’ or ‘high’ for that variable. For example, for ethnicity, the mean percent of the population which identified as white was averaged from the three counties. All chi-square tests were run using STATA 14.2. The result of an analysis was considered significant if p<0.05. Fischer’s Test was performed for tracts with less than five human cases of WNV.

### Logistic regressions

Logistic regressions were conducted to examine the relationship between the variables and the dependent dichotomous variable, which was the presence or absence of human WNV cases in each census tract. Analyses were conducted only for the three counties combined and were conducted using STATA 14.2. Environmental and socioeconomic variables used in analyses included temperature, precipitation, mosquitoes positive for WNV, habitat, age, education, median housing age, home vacancies, median income, population density, ethnicity, and language. Variables were tested to determine whether they were highly correlated. Minimum, mean, and max temperature were highly correlated, and only mean temp was used in the final model. Twelve variables were considered in the model. Most of the independent variables were continuous except for two variables, habitat and WNV positive mosquitoes, which were categorical. Habitat was treated as a dummy variable, with rural habitat represented by a ‘1’ and non-rural by a 0; WNV mosquitoes were similarly coded with census tracts having mosquitoes testing negative for WNV with a ‘0’ and those with mosquitoes testing positive for WNV with a ‘1’. The output was used to examine the overall chi-square value, and whether it was significant at *p*<0.05. A Z-score and its related p-value was determined for each variable in the model. Variables with Z-scores that had p-values <0.05 were considered significant. Finally, the odds ratios were determined for each variable in the model. For the three counties combined, three models were created: a model with only environmental factors, a model of only socioeconomic factors, and the final model with all twelve variables included. The odds ratios and 95% confidence intervals were reported for the adjusted associations estimated through these models.

## Results

### Human West Nile cases by county

The number of human cases of West Nile Virus (WNV) for 2011-2015 totaled 169 cases for the three counties (Table 1). The largest number of cases was in Stanislaus County (114), followed by San Joaquin County (39), and the lowest number of cases occurred in Merced County (16) (Table 1).

**Table 1.** The number of human West Nile virus cases in Merced, Stanislaus, and San Joaquin counties, from 2011-2015. Stanislaus has 94 total tracts; however, 93 tracts were used for the analysis. This lead to 281 (rather than 282) census tracts used.

| County | Total Human Cases 2011-2015 | No. of Census Tracts | Census Tracts with WNV | Census Tracts without WNV |
| --- | --- | --- | --- | --- |
| Merced | 16 | 49 | 10 | 39 |
| Stanislaus | 114 | 94 | 54 | 40 |
| San Joaquin | 39 | 139 | 33 | 106 |
| Total | 169 | 282 |  |  |

The WNV cases were plotted for each county for 2011-2015 (Fig 2). Stanislaus had the highest number of cases overall for all five years (Fig 2), followed by San Joaquin County, and Merced County had the lowest number of cases. A map with census tracts in each county was used to highlight the census tracts for presence or absence of human WNV cases (Figs 3–5). In San Joaquin County, there were 27 tracts with 1 human WNV case, and 5 tracts with 2 human cases, and 0 tracts with 3 or more cases (Fig 3). In Stanislaus County, there were 22 tracts with 1 case, 19 tracts with 2 cases, and 13 tracts with 3 or more cases (Fig 4). In Merced County, there were 7 tracts with 1 case, 1 tract with 2 cases, and 2 tracts with 3 or more cases (Figs 5,6).

**Fig 2.**
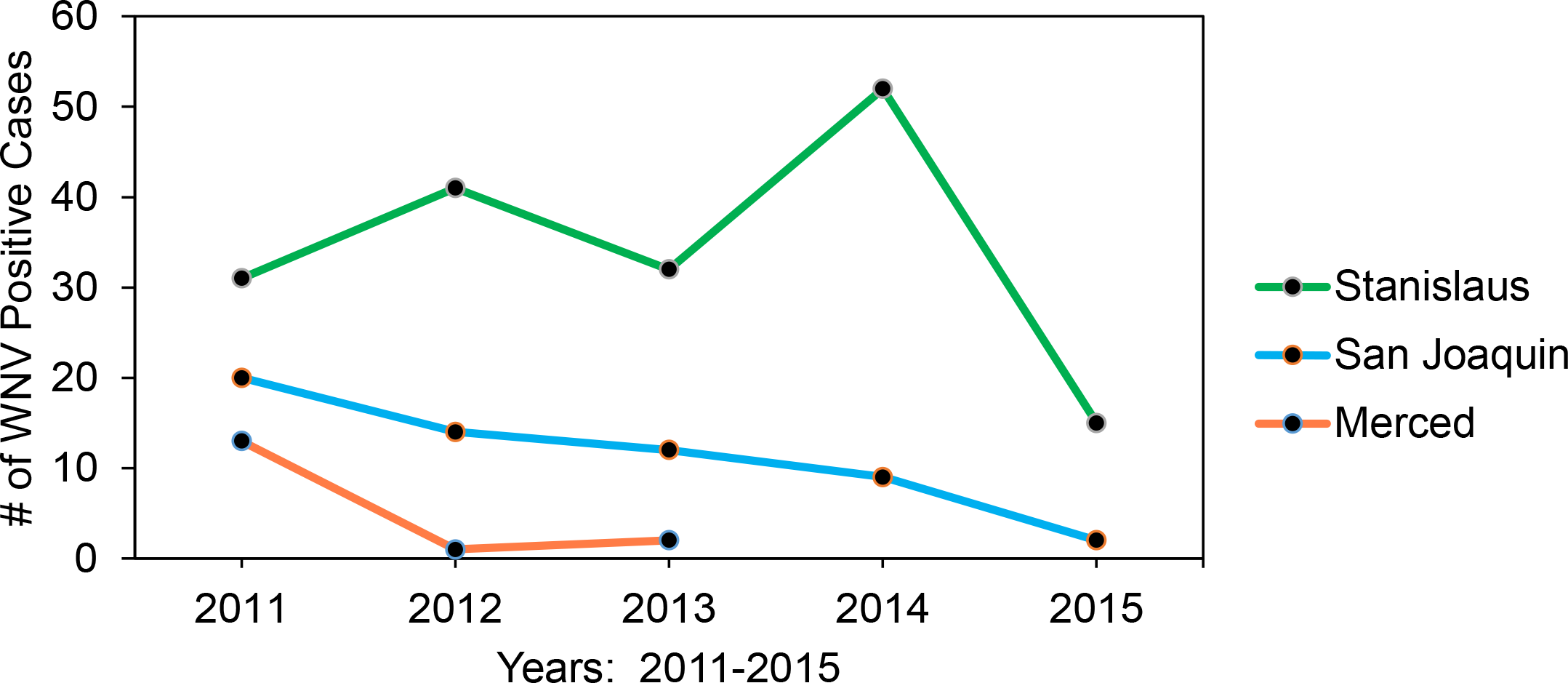
Human West Nile virus cases for San Joaquin, Stanislaus and Merced counties from 2011-2015.

**Fig 3.**
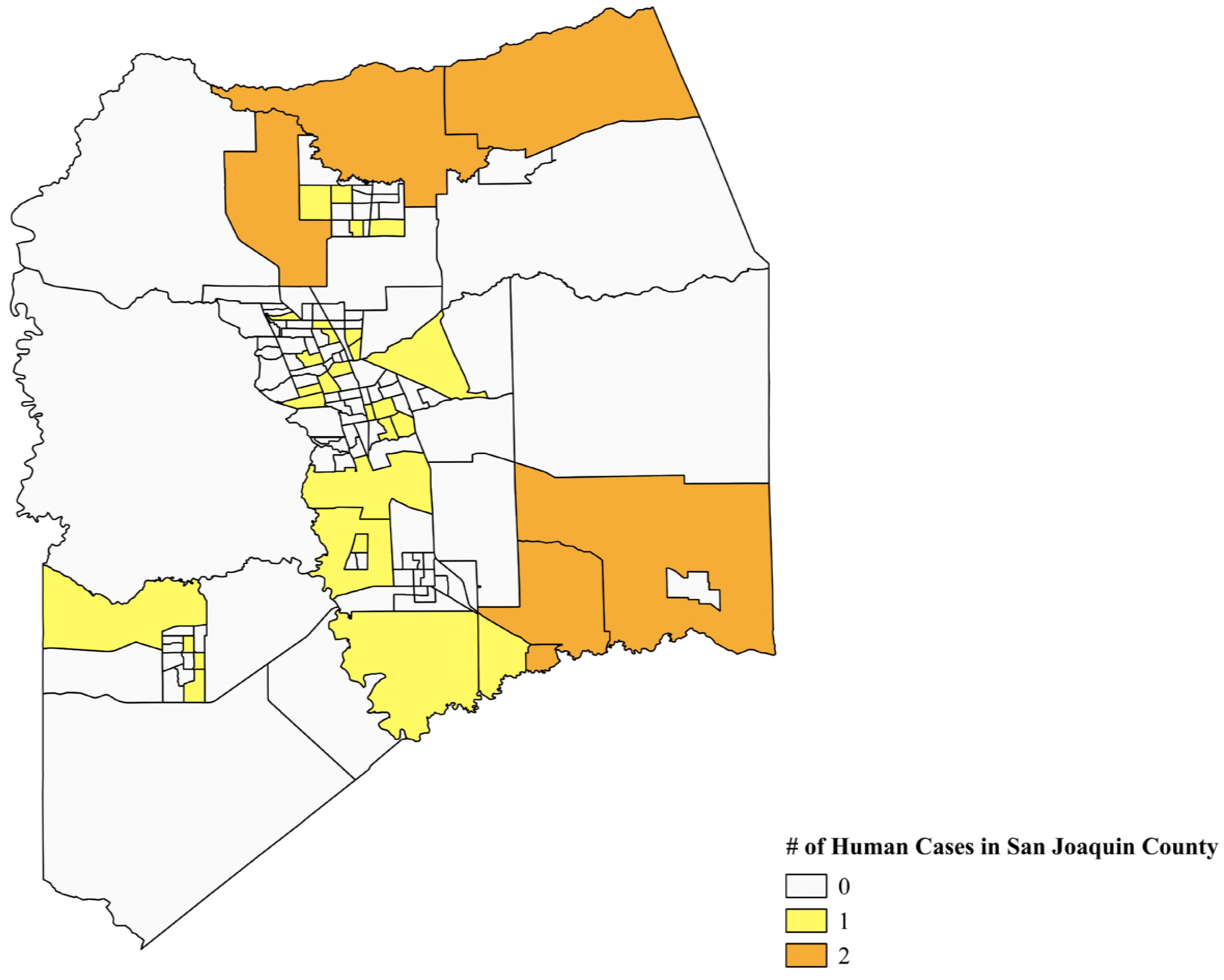
Map of San Joaquin County, California, census tracts. Tracts shaded in yellow have one human WNV case, while those in orange have 2 human cases. The map was made using the software QGIS 2.18.9 and census tract layers from the 2010 Census TIGER/Line files, accessed at https://qgis.org/en/site/ and https://www.census.gov/cgi-bin/geo/shapefiles2010/main.

**Fig 4.**
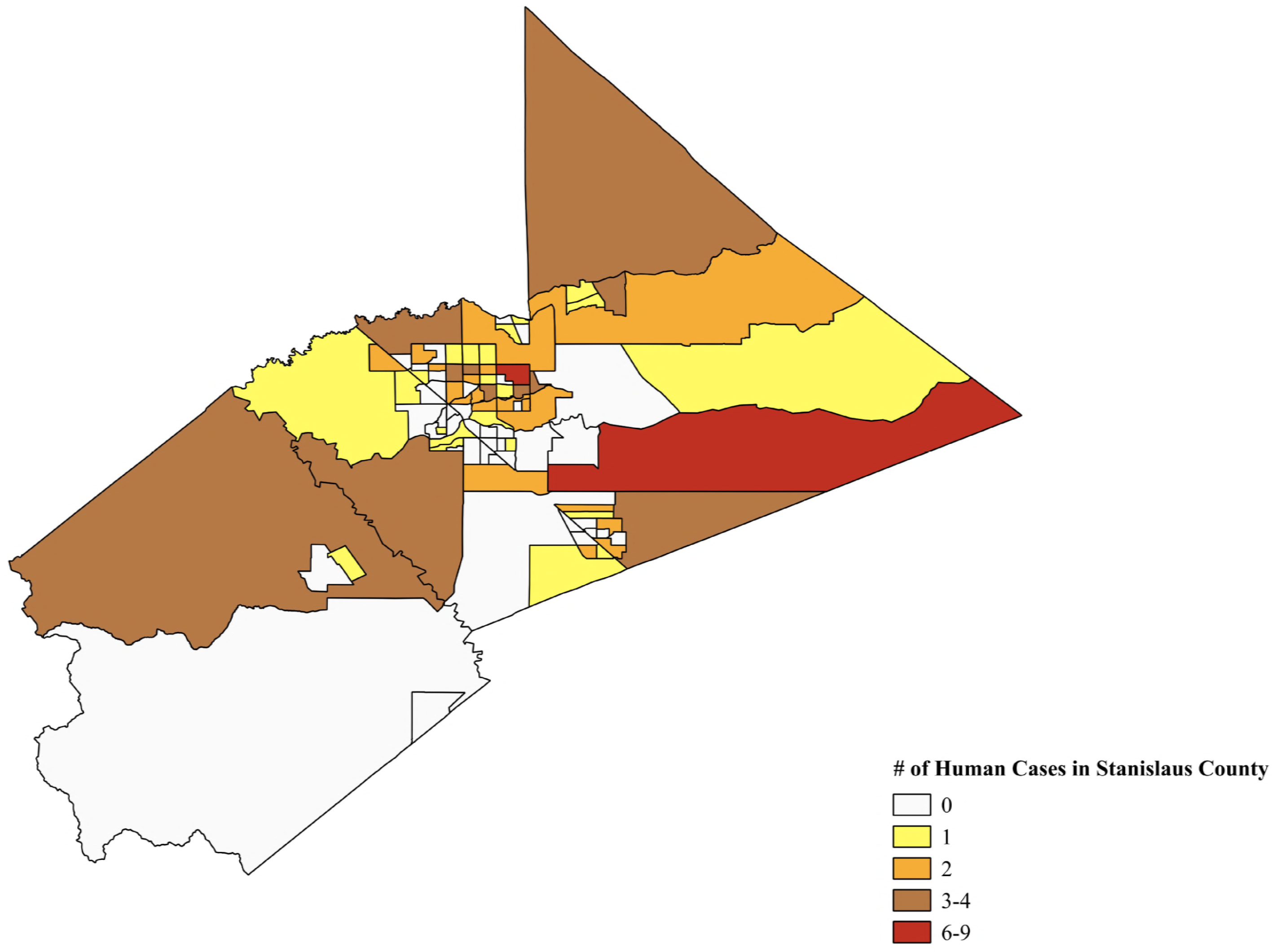
Map of Stanislaus County, California, census tracts. Tracts shaded in yellow have one human WNV case, while those in orange have 2 human cases. Tracts in brown have 3-4 cases, and tracts shaded in red have 6-9 human WNV cases. Map was made using software QGIS 2.18.9 and census tract layers from the 2010 Census TIGER/Line files, accessed at https://qgis.org/en/site/ and https://www.census.gov/cgi-bin/geo/shapefiles2010/main.

**Fig 5.**
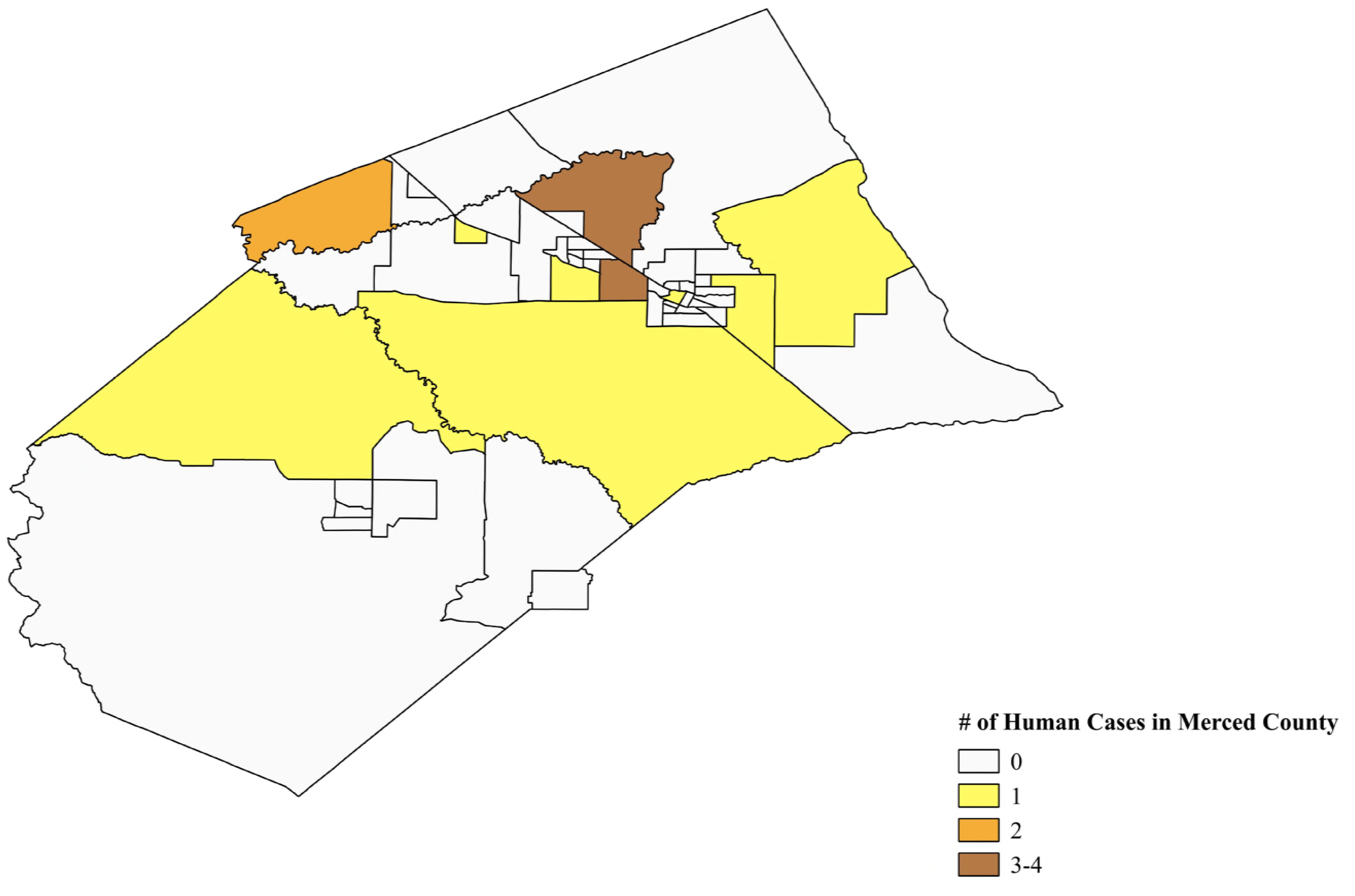
Map of Merced County, California, census tracts. Tracts shaded in yellow have one human WNV case, while those in orange have 2 human cases. Tracts in brown have 3-4 human WNV cases. Map was made using software QGIS 2.18.9 and census tract layers from the 2010 Census TIGER/Line files, accessed at https://qgis.org/en/site/ and https://www.census.gov/cgi-bin/geo/shapefiles2010/main.

**Fig 6.**
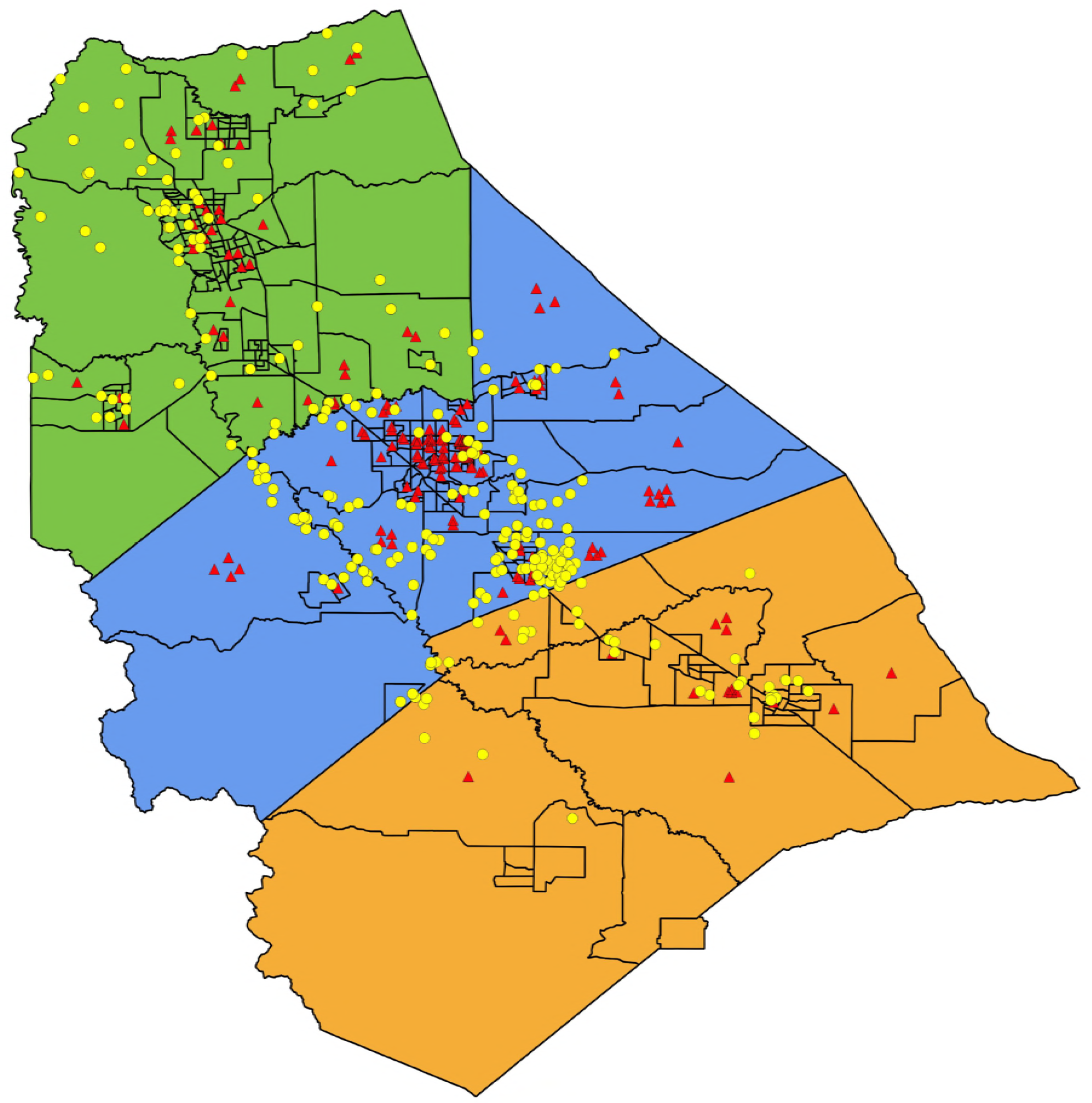
From north to south, San Joaquin, Stanislaus, and Merced counties. Yellow circles represent WNV positive mosquitoes (*Culex pipiens, Culex tarsalis*), while red triangles represent human cases of WNV. Map was made using software QGIS 2.18.9 and census tract layers from the 2010 Census TIGER/Line files, accessed at https://qgis.org/en/site/ and https://www.census.gov/cgi-bin/geo/shapefiles2010/main.

### Association tests for Factors and Human WNV cases

#### San Joaquin County

In San Joaquin County, 139 census tracts were used for chi-square tests, which included 39 human WNV cases (Table 1). A chi-square test of independence was performed for San Joaquin County to examine whether there was an association between WNV human cases and each of the ten variables previously mentioned. A significant relationship was found between human WNV cases and the following variables: tracts with high home vacancies, (*X*^2^_1_=4.709,*p* < .05, N=139), higher population density (*X*^2^_1_=5.832,*p* < .05, N=139), WNV positive mosquitoes, (*X*^2^_1_=11.29, *p* < .05, N=139) (Table 2), and finally habitat (*X*^2^_1_=4.993, *p* < .05, N=139) with more cases in urban tracts (tracts that were not rural; 24 non-rural vs. 9 rural). The relationship between age, education, median housing year, median household income, ethnicity, and language spoken at home and WNV cases was not significant (Table 2).

**Table 2.** Variables tested for association with human WNV cases in San Joaquin County. The mean value for each variable is shown. Census tracts with WNV were classified as ‘low’ or ‘high’ for each variable. The number of human WNV cases in tracts classified as low or high are shown. Chi-square value and p-value are shown; p-values were considered significant for P<0.05. *Mean is listed for all variables except income and housing year, where the US Census used median values.

| Variable | Mean* | Low | High | X<sup>2</sup> value/<br>p-value |
| --- | --- | --- | --- | --- |
| Positive WNV<br>Mosquito in tract or<br>not | | Human Cases<br>18 tracts | No human<br>cases<br>25 tracts | X<sup>2</sup>: 11.29<br>$P < 0.05^*$ |
| Habitat:<br>Rural or non-Rural | Rural:<br>Less than 2500<br>people per square<br>mile | Not rural<br>24 Cases | Rural<br>9 Cases | X<sup>2</sup>: 4.993<br>$P < 0.05^*$ |
| Age<br>65 and over | 65 and over:<br>10.4% | < 10.4%<br>17 Cases | ≥ 10.4%<br>16 Cases | X<sup>2</sup>: 0.001<br>$P > 0.05$ |
| Education:<br>High School<br>Graduate or Higher | 76.7% | < 76.7%<br>12 Cases | ≥ 76.7%<br>21 Cases | X<sup>2</sup>: 1.189<br>$P > 0.05$ |
| Median Housing<br>Year | 1979 | < 1979<br>17 Cases | ≥ 1979<br>16 Cases | X<sup>2</sup>: 0.264<br>$P > 0.05$ |
| Housing Vacancies | 32 units | < 32<br>10 Cases | ≥ 32<br>23 Cases | X<sup>2</sup>: 4.709<br>$P < 0.05^*$ |
| Median Household<br>Income | \$54,341 | < \$54,341<br>17 Cases | ≥ \$54,341<br>16 Cases | X<sup>2</sup>: 0.052<br>$P > 0.05$ |
| Population Density<br>(per sq. mile of<br>land area) | 492.6 | < 492.6<br>9 Cases | ≥ 492.6<br>24 Cases | X<sup>2</sup>: 5.832<br>$P < 0.05^*$ |
| Ethnicity: White or<br>Non-White | White: 51.0% | < 51.0%<br>15 Cases | ≥ 51.0%<br>18 Cases | X<sup>2</sup>: 0.10<br>$P > 0.05$ |
| Language:<br>English speaking | English: 61.4% | < 61.4%<br>15 Cases | ≥ 61.4%<br>18 Cases | X<sup>2</sup>: 0.006<br>$P > 0.05$ |

#### Stanislaus County

Chi-square independence tests were similarly calculated for Stanislaus County. In Stanislaus County, 94 census tracts were used for all chi-square tests except median housing year (which used 93 tracts), and 114 human WNV cases (Tables 1 and 3). The relationship between the following variables and WNV human cases was significant; tracts with higher than average household income were more likely to have WNV cases (*X*^2^_1_=4.921, *p* < .05, N=93) as were tracts with higher education levels (*X*^2^_1_=9.603, *p* < .05, N=93), and ethnicity was significant as well, with more WNV cases occurring in census tracts with a higher than average percent white residents (*X*^2^_1_=10.10, *p* < .05, N=93) (Table 3). In contrast, the relationship was not significant between WNV human cases and mosquitoes positive for WNV, habitat, population age, median housing year, home vacancies, population density and language (all *p* >.05, Table 3).

**Table 3.** Variables tested for association with human WNV cases in Stanislaus County. The mean value for each variable is shown. Census tracts with WNV were classified as ‘low’ or ‘high’ for each variable. The number of human WNV cases in tracts classified as low or high are shown. Chi-square value and p-value shown; p-values were considered significant for P<0.05. *Mean is listed for all variables except income and housing year, where the US Census used median values.

| Variable | Mean* | Low | High | X<sup>2</sup> value/<br>p-value |
| --- | --- | --- | --- | --- |
| Positive WNV Mosquito in tract or not | | Human Cases<br>23 tracts | No human cases<br>10 tracts | X<sup>2</sup>: 3.122<br>$P > 0.05$ |
| Habitat: Rural or non-Rural | Rural:<br>Less than 2500 people per square mile | Not rural<br>40 Cases | Rural<br>14 Cases | X<sup>2</sup>: 1.638<br>$P > 0.05$ |
| Age: 65 and over | 65 and over: 10.7% | < 10.7%<br>24 Cases | ≥ 10.7%<br>30 Cases | X<sup>2</sup>: 1.567<br>$P > 0.05$ |
| Education: High School Graduate or Higher | 75.3% | < 75.3%<br>19 Cases | ≥ 75.3%<br>35 Cases | X<sup>2</sup>: 9.603<br>$P < 0.05^*$ |
| Median Housing Year | 1979 | < 1979<br>27 Cases | ≥ 1979<br>26 Cases | X<sup>2</sup>: 0.022<br>$P > 0.05$ |
| Housing Vacancies | 33 | < 33<br>23 Cases | ≥ 33<br>31 Cases | X<sup>2</sup>: 0.906<br>$P > 0.05$ |
| Median Household Income | \$51,094 | < \$51,094<br>24 Cases | ≥ \$51,094<br>30 Cases | X<sup>2</sup>: 4.921<br>$P < 0.05^*$ |
| Population Density (per sq. mile of land area) | 344.2 | < 344.2<br>9 Cases | ≥ 344.2<br>45 Cases | X<sup>2</sup>: 1.734<br>$P > 0.05$ |
| Ethnicity: White or Non-White | White: 65.6% | < 65.6%<br>16 Cases | ≥ 65.6%<br>38 Cases | X<sup>2</sup>: 10.10<br>$P < 0.05^*$ |
| Language: English speaking | English: 59.8% | < 59.8%<br>19 Cases | ≥ 59.8%<br>35 Cases | X<sup>2</sup>: 1.447<br>$P > 0.05$ |

#### Merced County

In Merced County, 49 census tracts were included for analyses, and 16 human WNV cases (Tables 1 and 4). Chi-square independence tests found a significant association between history of WNV positive mosquitoes (*X*^2^_1_=9.70, *p* < .05, N=49) and WNV cases, but all other variables (habitat, age, education, housing age, home vacancies, income, population density, ethnicity, language) were not significantly associated with WNV human cases (Table 4).

**Table 4.** Variables tested for association with human WNV cases in Merced County. The mean value for each variable is shown. Census tracts with WNV were classified as ‘low’ or ‘high’ for each variable. The number of human WNV cases in tracts classified as low or high are shown. Chi-square value and p-value shown; p-values were considered significant for P<0.05. *Mean is listed for all variables except income and housing year, where the US Census used median values.

| Variable | Mean* | Low | High | X<sup>2</sup> value/<br>p-value |
| --- | --- | --- | --- | --- |
| Positive WNV<br>Mosquito in tract or<br>not | | Human<br>Cases<br>9 tracts | No human<br>cases<br>8 tracts | X<sup>2</sup>: 3.90<br>$P < 0.05^*$ |
| Habitat:<br>Rural or non-Rural | Rural:<br>Less than 2500<br>people per square<br>mile | Not rural<br>4 cases | Rural<br>6 cases | X<sup>2</sup>: 0.141<br>$P > 0.05$ |
| Age:<br>65 and over | 65 and over:<br>9.4% | < 9.4%<br>5 cases | ≥ 9.4%<br>5 cases | X<sup>2</sup>: 0.047<br>$P > 0.05$ |
| Education:<br>High School<br>Graduate or Higher | 67% | < 67%<br>4 cases | ≥ 67%<br>6 cases | X<sup>2</sup>: 0.291<br>$P > 0.05$ |
| Median Housing<br>Year | 1980 | < 1980<br>4 cases | ≥ 1980<br>6 cases | X<sup>2</sup>: 0.291<br>$P > 0.05$ |
| Housing Vacancies | 41 | < 41<br>3 Cases | ≥ 41<br>7 Cases | X<sup>2</sup>: 0.477<br>$P > 0.05$ |
| Median Household<br>Income | \$43,844 | < \$43,844<br>3 cases | ≥ \$43,844<br>7 cases | X<sup>2</sup>: 0.090<br>$P > 0.05$ |
| Population Density<br>(per sq. mile of land<br>area) | 132.2 | < 132.2<br>4 cases | ≥ 132.2<br>6 cases | X<sup>2</sup>: 0.201<br>$P > 0.05$ |
| Ethnicity: White or<br>Non-White | White: 58.0% | < 58%<br>3 cases | ≥ 58%<br>7 cases | X<sup>2</sup>: 0.720<br>$P > 0.05$ |
| Language:<br>English speaking | English:<br>48.5% | < 48.5%<br>2 cases | ≥ 48.5%<br>8 cases | X<sup>2</sup>: 0.278<br>$P > 0.05$ |

#### All three counties combined-association of WNV and factors

Data from San Joaquin, Stanislaus, and Merced County were pooled in order to conduct a combined chi-square test of independence for WNV human cases and their potential association with any of the ten previously mentioned variables (Table 5). Tracts where WNV positive mosquitoes were found were more likely to have WNV cases (*X*^2^_1_=23.06,*p* < .05, N=281). A significant relationship was found between WNV human cases and habitat (*X*^2^_1_ =7.199, *p* < .05, N=281) with urban tracts having 68 cases while 29 cases occurred in rural tracts (Table 5). Census tracts with higher than average levels of education had almost twice as many WNV cases than tracts with lower than average education levels (*X*^2^_1_=5.537,*p* < .05, N=281) (Table 5; 64 vs. 33 cases). Census tracts with higher home vacancies had higher human WNV cases (*X*^2^_1_=5.307,*p* < .05, N=281). Population density was significantly related to WNV cases, with higher density tracts having more cases of WNV (*X*^2^_1_=7.382,*p* < .05, N=281). Ethnicity was also significantly associated with human WNV cases with more WNV cases occurring in census tracts having a higher than average % white demographic (*X*^2^_1_=7.728, *p* < .05, N=281) (Table 5). Human cases of WNV were not associated with age (percentage 65 and over), median housing year, median household income, or language spoken at home (all *p* >0.05, Table 5).

**Table 5.** Variables tested for association with human WNV cases in the three counties combined. The mean value for each variable is shown. Census tracts with WNV were classified as ‘low’ or ‘high’ for each variable. The number of human WNV cases in tracts classified as low or high are shown. Chi-square value and p-value shown; p-values were considered significant for P<0.05. *Mean is listed for all variables except income and housing year, where the US Census used median values.

| Variable | Mean* | Low | High | X<sup>2</sup> value/<br>p-value |
| --- | --- | --- | --- | --- |
| Positive WNV Mosquito<br>in tract or not | | Human cases<br>50 tracts | No human<br>cases<br>43 tracts | X<sup>2</sup>: 23.06<br>$P < 0.05^*$ |
| Habitat:<br>Rural or non-Rural | Rural:<br>Less than<br>2500 people<br>per square<br>mile | Not Rural<br>68 cases | Rural<br>29 Cases | X<sup>2</sup>: 7.199<br>$P < 0.05^*$ |
| Age:<br>65 and over | 65 and over:<br>10.1% | < 10.1%<br>45 Cases | ≥ 10.1%<br>52 Cases | X<sup>2</sup>: 0.929<br>$P > 0.05$ |
| Education:<br>High School Graduate or<br>Higher | 73.0% | < 73.0%<br>33 Cases | ≥ 73.0%<br>64 Cases | X<sup>2</sup>: 5.537<br>$P < 0.05^*$ |
| Median Housing Year | 1979 | < 1979<br>48 Cases | ≥ 1979<br>48 Cases | X<sup>2</sup>: 0.983<br>$P > 0.05$ |
| Housing Vacancies | 35 | < 35<br>40 Cases | ≥ 35<br>57 Cases | X<sup>2</sup>: 5.307<br>$P < 0.05^*$ |
| Median Household<br>Income | \$47,469 | < \$47,469<br>39 Cases | ≥ \$47,469<br>58 Cases | X<sup>2</sup>: 2.067<br>$P > 0.05$ |
| Population Density<br>(per sq. mile of land<br>area) | 323.0 | < 323.0<br>23 Cases | ≥ 323.0<br>74 Cases | X<sup>2</sup>: 7.382<br>$P < 0.05^*$ |
| Ethnicity:<br>White or Non-White | White:<br>58.2% | < 58.2%<br>34 Cases | ≥ 58.2%<br>63 cases | X<sup>2</sup>: 7.728<br>$P < 0.05^*$ |
| Language: English<br>speaking | English:<br>56.6% | < 56.6%<br>34 Cases | ≥ 56.6%<br>63 Cases | X<sup>2</sup>: 2.260<br>$P > 0.05$ |

### Logistic regression

A multivariate analysis of only environmental factors found a positive association between human cases of WNV and precipitation (OR=1.4, *p*=0.003), mean temperature (OR=2.2, *p*=0.002), and WNV positive mosquitoes (OR=3.0, *p*=0.000) (Table 6). The logistic regression analysis between human cases of WNV and socioeconomic factors found ethnicity (OR=1.0, *p*=0.014) was statistically significant, while the other factors including age, education, housing age, housing vacancies, income, population density, and language were not (Table 7). Finally, the combined logistic regression analysis of all 12 variables found that precipitation (OR=1.5, *p*=0.005), mean temperature (OR=2.8, *p*=0.002), and WNV positive mosquitoes (OR=3.1, *p*=0.000) were statistically significant (Table 8).

**Table 6.**
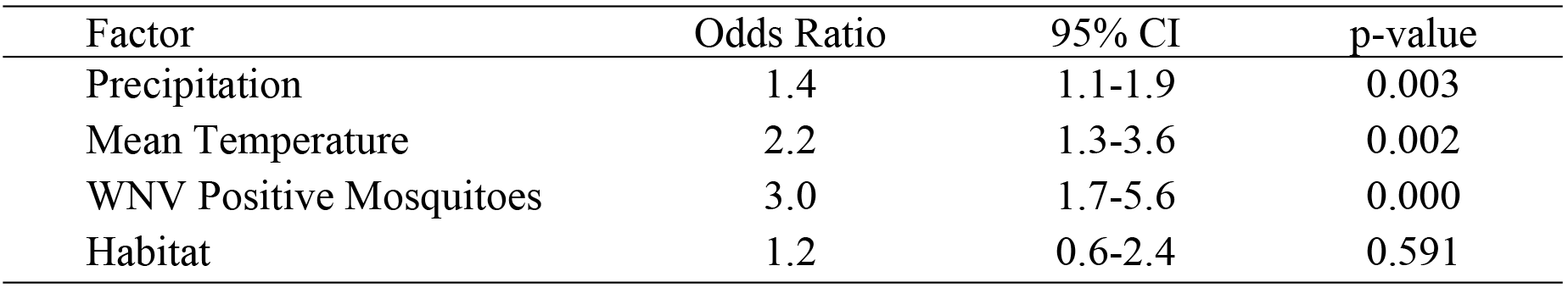
Logistic regression for the three counties combined, including only environmental factors; odds ratios and 95% confidence intervals (*X*^2^_1_=42.18, *p* < .0001).

| Factor | Odds Ratio | 95% CI | p-value |
| --- | --- | --- | --- |
| Precipitation | 1.4 | 1.1-1.9 | 0.003 |
| Mean Temperature | 2.2 | 1.3-3.6 | 0.002 |
| WNV Positive Mosquitoes | 3.0 | 1.7-5.6 | 0.000 |
| Habitat | 1.2 | 0.6-2.4 | 0.591 |

**Table 7.** Logistic regression for the three counties combined, including only socioeconomic factors; odds ratios and 95% confidence intervals (*X*^2^_1_=20.69, *p* = 008).

| Factor | Odds Ratio | 95 % CI | p-value |
| --- | --- | --- | --- |
| Age | 1.0 | 0.9-1.0 | 0.825 |
| Education | 1.0 | 0.9-1.0 | 0.587 |
| Housing Age | 0.9 | 0.9-1.0 | 0.902 |
| Housing Vacancies | 1.0 | 0.9-1.0 | 0.353 |
| Income | 1.0 | 0.9-1.0 | 0.585 |
| Population Density | 0.9 | 0.9-1.0 | 0.371 |
| Ethnicity | 1.0 | 1.0-1.0 | 0.014 |
| Language | 0.9 | 0.9-1.0 | 0.352 |

**Table 8.** Logistic regression for the three counties, including all 12 factors; odds ratios and 95% confidence intervals (*X*^2^_1_=48.43, *p* < .0001).

| Variable | Odds Ratio | 95 % CI | p-value |
| --- | --- | --- | --- |
| Precipitation | 1.5 | 1.1-2.1 | 0.005 |
| Mean Temperature | 2.8 | 1.4-5.7 | 0.002 |
| WNV Positive Mosquitoes | 3.1 | 1.6-6.2 | 0.001 |
| Habitat | 1.5 | 0.6-3.7 | 0.346 |
| Age | 1.0 | 0.9-1.1 | 0.233 |
| Education | 1.0 | 0.9-1.0 | 0.503 |
| Housing Age | 0.9 | 0.9-1.0 | 0.171 |
| Housing Vacancies | 1.0 | 0.9-1.0 | 0.346 |
| Income | 1.0 | 0.9-1.0 | 0.264 |
| Population Density | 1.0 | 0.9-1.0 | 0.385 |
| Ethnicity | 0.9 | 0.9-1.0 | 0.695 |
| Language | 0.9 | 0.9-1.0 | 0.424 |

## Discussion

In this study, we examined the influence of factors on the WNV human cases reported in three counties in the Central Valley of California. WNV incidence in humans was associated with numerous factors. Key factors that contributed to human cases included the environmental factors precipitation and temperature, along with the presence of West Nile positive mosquitoes; this was consistent with findings of previous studies. Socioeconomic factors which were found to be significant varied by county, and included increasing population density and home vacancies, along with higher income and education. The majority of studies examining environmental and socioeconomic factors have been conducted in urbanized areas (11,18,30). Rural areas have been studied to a lesser degree in terms of research related to factors associated with WNV.

In this study, the environmental factors temperature and precipitation were significantly related to the incidence of human West Nile cases, and this finding is consistent with previous studies (12,14,31,32). Mosquitoes need water for larvae to develop, and insects grow more rapidly in warmer temperatures (33–35). Standing water such as puddles, in gutters and storm drains, and habitats such as abandoned or neglected pools are ideal habitats for mosquitoes such as *Cx. pipiens*, a primary vector associated with WNV (19). There was a significant association overall of vacant homes and human cases of WNV. This is not surprising, since vacant homes may have neglected swimming pools which can serve as breeding grounds for mosquitoes which vector WNV such as *Culex* spp. (19). Along with temperature and precipitation, the presence of WNV positive mosquitoes was a strong predictor of human WNV cases. These findings are highly supportive of mosquito abatement efforts, whose control programs often focus on reducing mosquito populations where there are favorable habitats for their development. When environmental factors were considered separately or together with socioeconomic factors in logistic regression models, environmental factors were the most significant predictors associated with WNV cases in the region of this study.

For each of the three counties, the significant socioeconomic factors associated with the prevalence of WNV varied. For example, in San Joaquin County, a significant association was found between human cases of WNV and urban areas, high population density, and housing vacancies. In Stanislaus County, WNV cases were significantly associated with higher education, higher median household income, and increasing percent white population in census tracts. In Merced County, none of the socioeconomic factors were significantly associated with the presence of WNV cases. However, Merced had a small number of reported WNV cases (16) over the five years included in the study. For the three counties combined, human cases of WNV were associated with urban tracts, housing vacancies, increased population density, education, increasing percent Caucasian ethnicity, and English language speaking populations.

Previous studies have found an association between urbanization and increased WNV activity (11,18,21,36). Although the Northern San Joaquin region of the Central Valley is a rural area overall, areas in the region with increasing population density were significantly associated with human cases of WNV, as was non-rural (i.e. urban) habitat. In studies of urban areas, higher education levels were found to be significantly related to WNV human cases (21), which was similar to the finding of this study. Specifically, increasing levels of high school graduates or those with higher education were significantly associated with human cases of WNV. It is possible that residents in areas who have attained a higher education level are more likely to be informed of WNV and request WNV testing from their physicians. Investigators in Chicago and Detroit found that middle class neighborhoods were associated with high WNV activity and human infection (37). In contrast, the risk of WNV human infection was associated with low income areas in Orange County, CA, Harris County, TX, and in Shelby County, TN (15,18,38,32). Although in this study the three-county analysis did not find a significant relation of WNV with income, in one of the counties (Stanislaus County) which had the highest number of human WNV cases (114), higher income was significantly associated with WNV.

Ethnicity was significantly related to WNV cases in the overall three-county analysis, as well as being significant for Stanislaus County; census tracts with higher than average percent white residents were associated with human WNV cases. This study along with those previously mentioned suggest that socioeconomic factors including ethnicity, income, and education are closely intertwined with human risk of WNV. Wealthier neighborhoods may provide a favorable habitat for mosquitoes consisting of more open space than in lower income areas, and may have a certain combination of landscape vegetation that is preferable for mosquito breeding; sprinklers for watering vegetation may be more abundant as well. Overall, these results show that WNV human cases were significantly associated with a high percentage of people who are Caucasian, high school graduates or of higher education, and have high income. We suspect that the increased number of cases in Stanislaus County is not because there are more mosquitoes or more cases of WNV in the area. We believe that a possible explanation for these cases is that residents in higher income areas may be more likely to visit their doctor, request WNV testing from their doctor, or be willing to pay for the WNV testing.

WNV disease is a nationally notifiable condition; however, many cases are underreported as the majority of infections are asymptomatic (10,39). Under the physician’s care, a patient may be tested if they are showing symptoms of WNV and results are reported to local public health authorities if there is a positive case. We suspect that in most of these reported cases, patients are of high income and education and are aware of symptoms associated with WNV, therefore they request WNV testing. Testing must be done for WNV to confirm the disease and to differentiate it from other vector-borne (arboviral) diseases. This in turn means people of low socioeconomic status may not be aware that WNV is active in their area or what symptoms are associated with this disease. This lack of information prevents them from requesting WNV testing from their physician. It is imperative that members of the community are educated about prevention of vector-borne disease, and how to decrease their risk of being bitten and infected. This will require providing more educational materials to the community and depending on the area, providing them in the different languages spoken in the region.

Language spoken at home was not a significant factor associated with WNV cases in this study. Prior to this study, we suspected that the Hispanic population may have a language barrier in receiving WNV notices or health advisories, but that was not apparent in the results. It is possible that socioeconomics, for example lower income residents without insurance or less education, are unaware of symptoms of WNV and may be less likely to visit a doctor and ask for WNV testing (30). This could contribute to the finding of why WNV cases were associated with this combination of social factors: Caucasian, high school graduate or higher, and high income. This interrelatedness of social factors is complex and requires further investigation. In addition, educational materials on prevention of WNV, symptoms of the disease, and testing and treatment, could be targeted at lower income areas in the region. Although language did not appear as a significant factor in this study, it is possible that educational materials and programs in additional languages spoken in the area (Spanish, Hmong, Punjabi) would increase awareness and testing for WNV. Awareness of vector-borne disease in this region of California is timely, as more invasive species of mosquitoes capable of transmitting additional vector-borne diseases are introduced into the area.

It has been previously reported that with advancing age, the incidence of neuroinvasive disease and death due to WNV increases (40). However, results in this study showed that census tracts with more people ages 65 or over were not significantly associated with WNV cases for either San Joaquin, Stanislaus, Merced, or the three counties combined. Although the factor age (65 and over) was not significantly associated with human cases in this study, people of all ages remain susceptible to infection and surveillance should continue to target areas with all age groups. Other studies found housing age was strongly associated with human cases of WNV, yet it was not strongly associated with WNV cases in the analysis of the three counties individually or combined in this study. Housing age has been a significant factor in prior studies as a result of old storm water sewer systems that can serve as a breeding source for *Culex* mosquitoes (20,41). Consideration of storm drain types in housing tracts of different ages merits consideration when considering control programs. As mentioned previously, environmental factors were most strongly associated with the presence of WNV cases in the region of this study.

This analysis of environmental and socioeconomic factors that influence human cases of WNV can contribute to mosquito abatement districts’ efforts towards reducing mosquito populations and containing these risk factors associated with WNV. Education and communication are key elements in public health prevention programs. Both are of key importance during a disease outbreak in order to adequately inform citizens of the risk of WNV and how to protect themselves. It is critically important that prevention information is available to all populations represented in each county. Public health prevention messages could be created in other languages to inform the public especially those at greater risk of the dangers of WNV. Prevention messages such as “Drain standing water”, “Use window screens” “Mosquito proof your home”, “Apply insect repellent”, “Avoid mosquito bites”, and “West Nile virus is found throughout California” may be used. Mosquito abatement districts can expand their form of communication to the general public through pamphlets, newspaper ads, TV advertisements, billboards, and town hall meetings. Greater involvement with social media, a platform that mosquito abatement districts utilize, is needed to inform the greater public and could be used in multiple languages to reach all ethnic groups living in this region of the Central Valley. Health education needs to be continuous to help members of the community to stay active in protecting themselves against WNV and other emerging vector-borne disease in the area.

## Acknowledgements

We thank the Merced County Mosquito Abatement District (MCMAD), Alan Inman, Jason Bakken, Arlilla Bueno, and Rhiannon Jones of MCMAD for feedback during the project. University of California Merced undergraduate laboratory assistants Karen Cedano, Andrew Loera, and Lindsay Robson assisted with various aspects of the study. David Heft and Monica Patterson of Turlock Mosquito Abatement District provided helpful discussions.

## Supporting Information

**S1 Table.** Variables used in Stata Analyses.

| **Variable** | **Definition** |
| --- | --- |
| Precipitation | Annual precipitation data for the year 2011 was acquired from the Oregon State University Prism Climate Group. |
| Temperature | Mean temperature for 2011 was acquired from the Oregon State University PRISM Climate Group. |
| Mosquitoes<br>Pos.WNV | Positive mosquitoes for WNV in a tract. |
| Habitat | Using coordinates of census tracts and google earth, the tract's habitat was categorized as either rural or urban. |
| Age | Age group: 65 years and over were retrieved from the U.S. Census Bureau. |
| Education | The education category "Percent high school graduates or higher" was used. This information was retrieved from the U.S. Census Bureau. |
| Housing Age | Year structure built refers to when the building was first constructed. This information was retrieved from the U.S. Census Bureau. |
| Household<br>Vacancies | The category 'housing vacancies-other' was the number of home foreclosures according to the U.S. Census. |
| Income | Median income divides the income distribution into two equal groups, one having incomes above the median, and other having incomes below the median. The Census Bureau collected this data from surveys such as the American Community Survey (ACS). |
| Population<br>Density | Population density was retrieved from the U.S. Census Bureau. It is the population density per square mile of land area. |
| Ethnicity | Percentages of White ethnicity within a tract were acquired from the U.S. Census Bureau |
| Language | The percent speaking only English at home was retrieved from the U.S. Census Bureau. |

